# Dynamic Extreme Aneuploidy (DEA) in the vegetable pathogen *Phytophthora capsici* sheds light on instant evolution and intractability

**DOI:** 10.1101/297788

**Authors:** Jian Hu, Sandesh Shrestha, Yuxin Zhou, Xili Liu, Kurt Lamour

## Abstract

Oomycete plant pathogens are notoriously difficult to control, and individual isolates are highly unstable; making routine research challenging. Sequencing reveals extreme aneuploidy for single-spore progeny of the vegetable pathogen *Phytophthora capsici*; a phenomenon dubbed Dynamic Extreme Aneuploidy (DEA). Although extreme, the aneuploidy appears to be moderately stable. A single sporulating plant lesion may produce an armada of genetically unique individuals and helps explain the rapid increase of advantageous alleles (e.g. drug resistance), mating type switches to allow sex and the widely observed phenomenon, loss of heterozygosity (LOH). Investigation of other oomycetes indicate this phenomenon is not unique to *P. capsici*.

## Introductory

Although oomycete plant pathogens (water molds) look and act like fungi, they are distantly related and most antifungal compounds have no effect ^1^. Oomycetes are notoriously difficult to control ^2–4^. In research, isolates are highly unstable; making routine research challenging ^5,6^. Here we show that ploidy amongst the chromosomes in an individual isolate varies dramatically for asexual spores of *Phytophthora capsici*; a phenomenon dubbed Dynamic Extreme Aneuploidy (DEA). The number of individual chromosomes allocated to a single zoospore is highly variable and appears moderately stable. The practical consequence is that infected plants can release an armada of genomically unique individuals – responding quickly to selection pressures. DEA explains the rapid increase of advantageous alleles (e.g. drug resistance)^7^, mating type switches to allow sex and loss of heterozygosity (LOH) ^8–10^ and may spur novel investigation of yeast and human tumor tissues where loss of heterozygosity is reported^11,12^.

The genus Phytophthora has many destructive and pervasive plant pathogens ^2^. The 120+ species attack almost all dicot plants and threaten ecosystems and entire plant industries ^2,4^. For more than 150 years, scientists have struggled to manage this unwieldy genus and make of it (or, at least one of its members) a model organism ^13^. Part of this quest has focused on the vegetable pathogen, *P. capsici* – a devastating pathogen of vegetables ^13–15^. Ostensibly, an ideal research model; isolates can be immortalized under liquid nitrogen, grow rapidly on simple media, can be easy to mate in the laboratory, are often highly pathogenic and make copious spores with little coaxing. The problem is - sometimes an isolate will do all the above and sometimes it won’t. The intractability of Phytophthora (or success, depending on your point of view) lies in its plasticity ^5^. Phytophthora, when needed most (e.g. for a long-term laboratory, greenhouse or field experiments), is entirely unreliable.

The ploidy of oomycetes was debated actively for over 75 years and it wasn’t until Eva Sansome, in a 1961 Letter to Nature, provided convincing photomicrographic evidence of meiosis with *Pythium debaryanum* Hesse that the issue approached resolution ^16^. She suggested *P. debaryanum* was not unique and subsequent work proved her correct ^2,17^. She states “No metaphases were seen in the hyphae or sporangia, even after camphor treatment, and it is concluded that vegetative nuclei do not normally pass through an ordinary metaphase stage when dividing.” She concludes this brief letter stating “… members of this group may be diploid rather than haploid as has been previously supposed.” Her findings, although compelling, were not immediately accepted as many researchers recorded aberrant sexual and asexual inheritance patterns ^18, 5^.

In 2005, the NSF and USDA jointly funded a high quality whole genome sequence and a single nucleotide polymorphism (SNP) resource for *P. capsici*. With a strong team of scientists, money and motivation; a quality genome, ready for publication, was expected in months. Parents and what appeared to be normal progeny had been produced previously and a dense genetic map – a crucial tool missing from the Phytophthora research inventory was planned. Events did not unfold as expected and months turned into years. *Phytophthora capsici* carries a significant load of polymorphism in the form of heterozygous SNPs, often 1 every 100bp within a single genome ^8^. Initially, the construction of a genetic map proved impossible as many loci had aberrant inheritance patterns. Finally, it was discovered that short and long tracts (300bp to 1Mbp) of the parent and progenies genomes had spontaneously switched (mitotically) to one or the other parental haplotype – a phenomenon known as copy neutral Loss of Heterozygosity (LOH) ^9^. Removal of the loci residing in LOH tracts helped produce a detailed molecular map and the annotated genome were published in 2012 ^8^. Population studies using SNP markers reveal LOH is common in *P. capsici* and other oomycetes at locations worldwide ^10,19–22^.

Until recently, the 917 genetic scaffolds were not arranged according to the chromosomes (18 linkage groups) ^8^. In 2017, the map-ordered scaffolds were used to visualize heterozygous allele frequencies for *P. capsici* recovered from pepper in Taiwan and isolates of the closely related pathogen of taro, *P. colocasiae*, from Hawaii, Vietnam, China and Nepal ^23,24^. Whole genome and targeted-sequencing revealed ploidy varied by chromosome within individual isolates. Here we present novel data from single-zoospore isolates of *P. capsici* that further characterize this phenomenon including the distribution of heterozygous SNP allele frequencies, mating type stability and variation in sensitivity to the phytophthora-toxic chemical mefenoxam. These findings, in total, indicate the partitioning of chromosomes during mitotic zoospore development is asynchronous and likely plays an important role in adaptation and evolution.

Without the scaffold-ordered genetic linkage map, it was impossible to visualize chromosome-level variations in copy number. Analyses using unordered scaffolds can be deceptive ^25,26^. For example, visualization of heterozygous allele frequencies using the 917 un-ordered scaffolds from the *P. capsici* reference genome for a field-isolated parent and single-zoospore progeny suggest the genomes are primarily diploid or triploid (Figure 1). The same data, visualized by chromosome, indicates ploidy of the larger chromosomes obscures the allele distributions for the smaller chromosomes (Figure 2). Linkage group 9 provides an exquisite example where the parent and four mitotic offspring vary between the di, tri, hexa and tetraploid states (Figure 2). Analysis of whole genome data for 33 isolates of *P. capsici* (or near relatives such as *P. tropicalis* and *P. colocasiae*) from locations worldwide indicate less than half exist at a single, homogenous, level of ploidy (Supplementary Figures).

**Figure 1.**
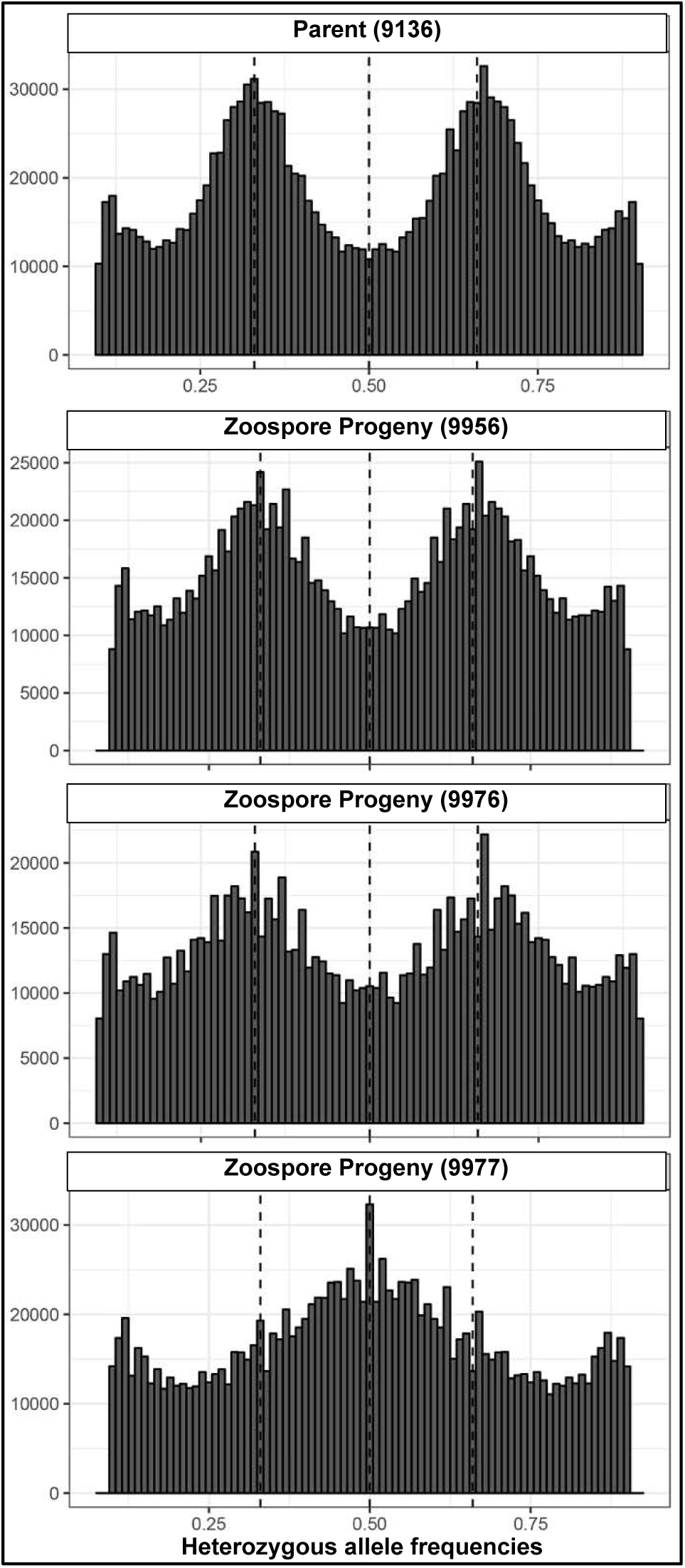
Histograms showing whole genome heterozygous allele frequencies for parent and zoospore progeny of the vegetable pathogen, *P. capsici*, based on mapping to the unordered 917 scaffolds of the *P. capsici* reference genome. Dotted lines indicate 25, 50 and 75% allele frequencies.

**Figure 2.**
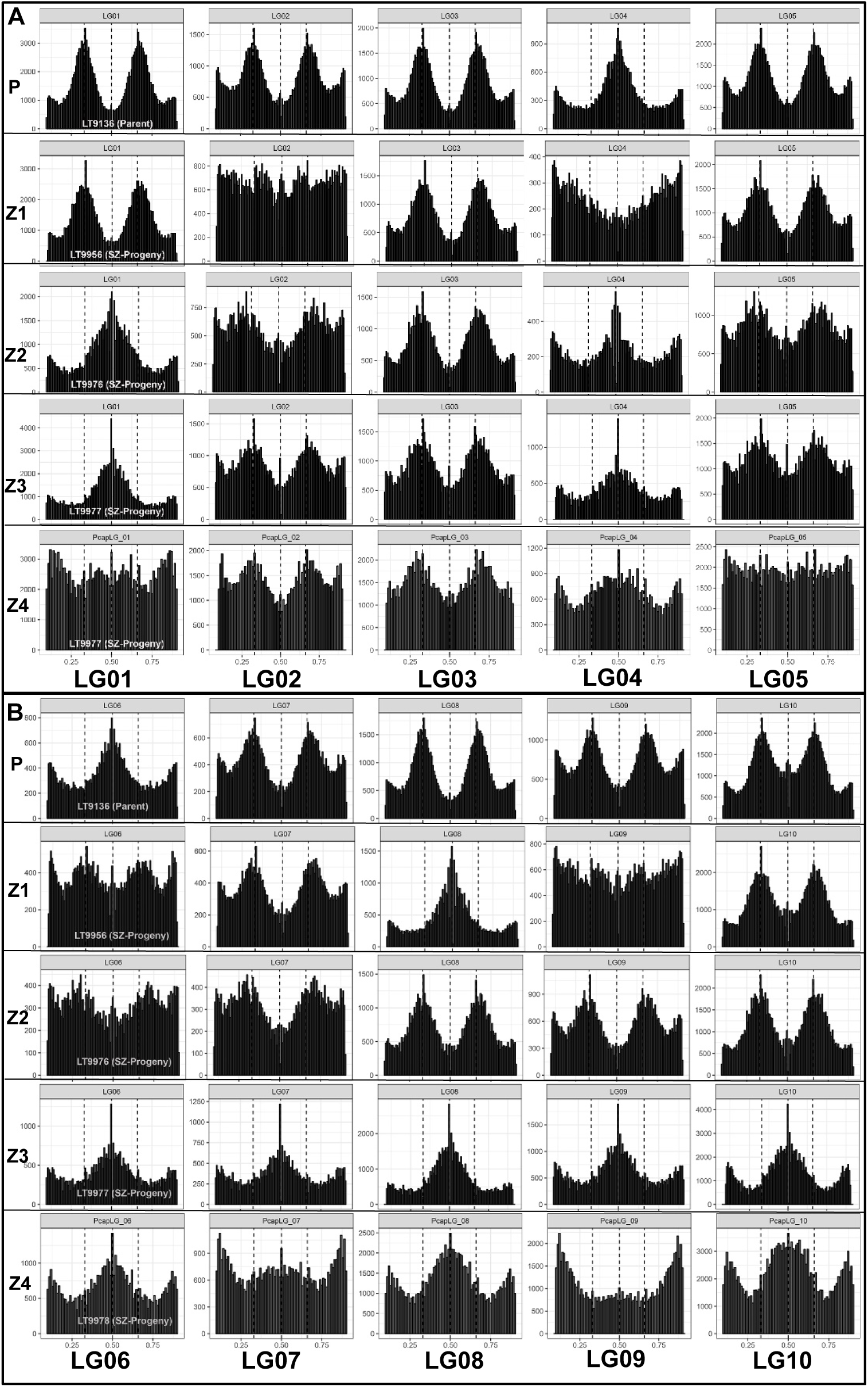
Linkage-group specific ploidy for a field isolate of *Phytophthora capsici* (P) and four mitotic zoospore progenies (Z1 to Z4). Each row shows histograms of heterozygous allele frequencies for parent and progeny isolates across the ten largest linkage groups. The y-axis denotes the number of markers and the x-axis their allele frequencies with dotted vertical lines at 33, 50 and 66%. A, linkage groups 1 to 5 and B, linkage groups 6 to 10.

**Figure 2.**
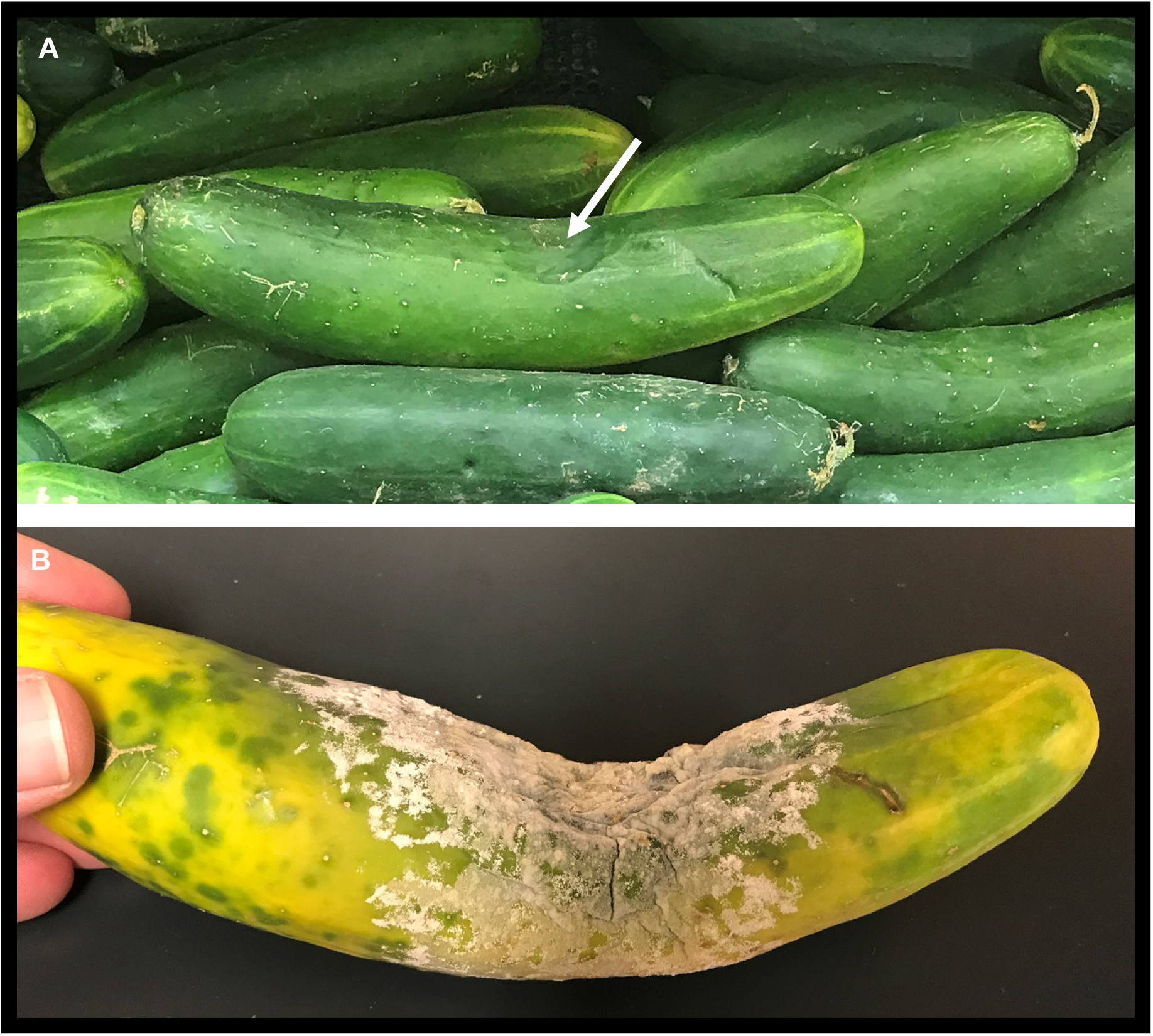
Cucumber fruit recovered August 3, 2017 from major supermarket chain in Knoxville, Tennessee (origin unknown). **A**, white arrow designates sunken, firm lesion on specific fruit at the store. **B**, the same fruit producing millions of asexual sporangia-spores after three days incubation under ambient (outdoor) conditions (K. Lamour, Knoxville, TN, Aug 3-6). Immersion of the whole fruit in water produced >100 million bi-flagellate swimming zoospores.

An important implication for *P. capsici* (and likely other outcrossing oomycetes) is the establishment of invasive long-lived populations through mating type switches. *Phytophthora capsici* requires the interaction of an A1 and A2 mating type to produce thick-walled, long-lived sexual dormancy spores (oospores) ^2^. *Phytophthora capsici* is an invasive pathogen in the US ^27^. Yet, newly introduced populations, even those emanating from a small focus of initial infection (e.g. a single infected plant), often accomplish sexual outcrossing within the first year of an epidemic ^27^. Analysis of 384 single-zoospore isolates, derived from three A1 and three A2 mating type isolates indicate mating type is highly unstable for A2 isolates. All 140 of the A1-derived isolates remained true to the A1 mating type. For the 241 A2-derived isolates, only 26% remained A2, 11% switched to A1 and 63% became self-fertile. Recently, genetic analysis of a bi-parental inbreeding field population of *P. capsici* revealed the A2 mating type requires elevated heterozygosity across a distinct mating-type region and DEA may play an important role in the development of sexually-active populations, even where the introductory event was limited to a single A2 mating type. ^28^.

Mefenoxam is a commonly used oomycete-toxic compound with a site-specific mode of action and resistance occurs rapidly ^2,7^. In *P. capsici*, resistance is mediated by co-dominant loci of major effect, although the exact mutations are unknown ^7^. In Michigan, close monitoring of field populations exposed to high doses of mefenoxam revealed rapid acquisition of full resistance without a concomitant genetic bottleneck ^27^. Fully resistant populations appeared to be as genetically diverse as nearby sensitive populations suggesting the resistance allele somehow bypassed the two rounds of outcrossing (and mandatory dormancy) needed for meiosis to produce homozygous (fully resistant) individuals ^29^. Here, we tested 254 single-zoospore isolates from sensitive and intermediately sensitive field isolates and found progeny from sensitive isolates almost always remained sensitive and progeny from intermediately sensitive isolates ranged from intermediate to fully resistant – most likely driven by the increased copy number of the resistance allele that occurs via DEA (Figure 3).

**Figure 3.**
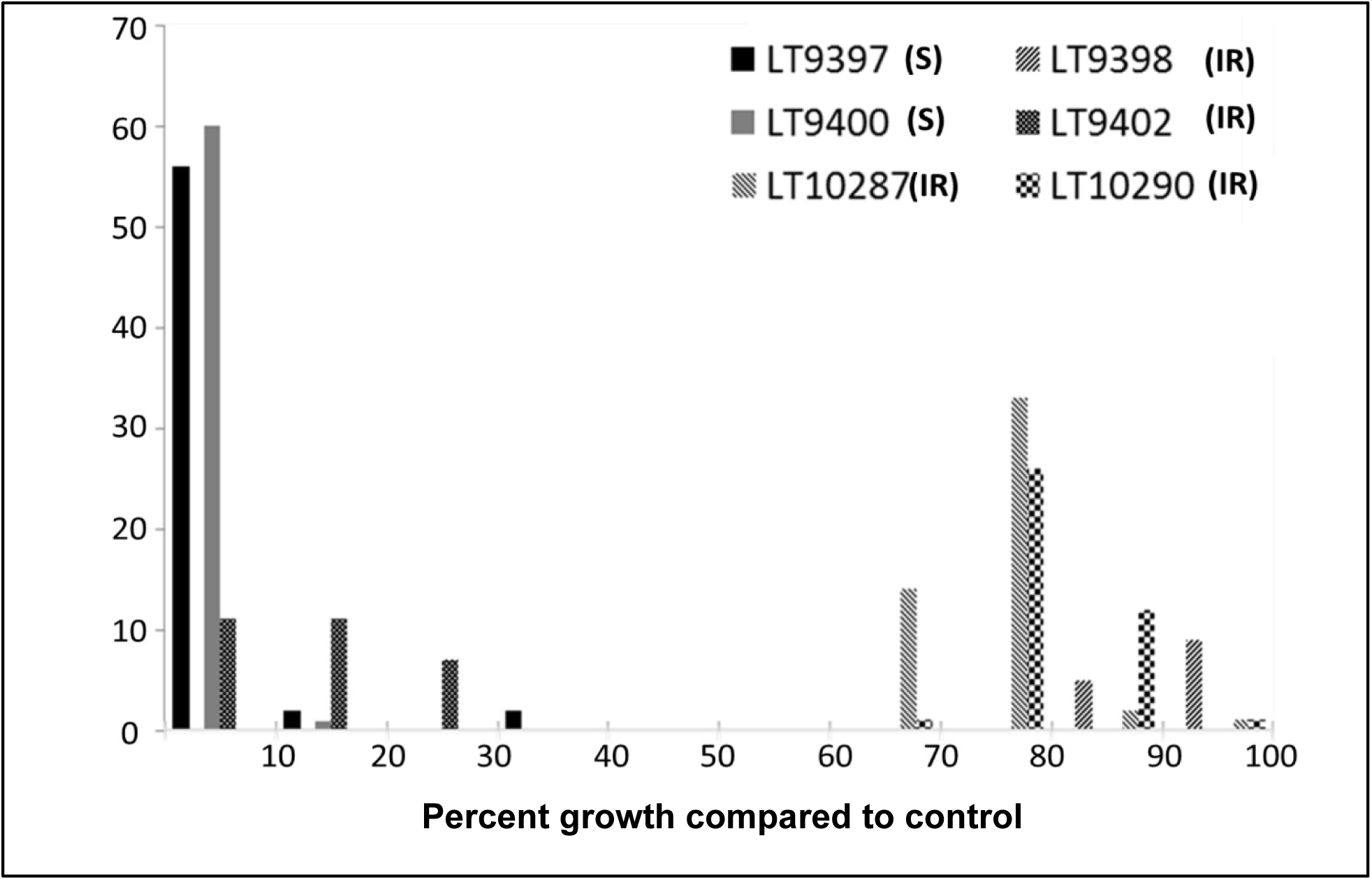
Response of 254 *Phytophthora capsici* single zoospore progeny from six field isolates to the oomycete-toxic chemical, mefenoxam. The six parental field isolates are listed in the upper right and denoted with various fill patterns. Parental isolates are either sensitive (S) to mefenoxam where their ability to grow on mefenoxam media is low compared to growth on un-amended control media or intermediately resistant (IR) where they can grow at roughly half their normal rate of growth on amended media. The progeny, charted as a histogram, ranges from fully sensitive to fully resistant.

To underscore the latent evolutionary potential within a single isolate, we recently assessed asexual zoospore production for a cucumber fruit on sale at a Knoxville, TN supermarket (August, 2017) (Figure 4). A fruit with a distinctive sunken lesion, typical for *P. capsici*, was incubated under ambient conditions (warm and humid) for three days and then immersed in 500ml of sterile water for one hour to stimulate release of the swimming zoospores. Hemocytometer counts indicate >100 million swimming zoospores were released. Zoospores can swim for days, are negatively geotropic (swim up) and chemotactic (swim towards plant exudates) ^2^. If *P. capsici* adhered strictly to the rules of mitosis, each zoospore should be an identical copy of the parent genome. Population studies using single-zoospore, hyphal-tip and/or infected-plant derived genomic DNA reveal this is not the case ^4^. Field populations, especially during the explosive asexual phase of an epidemic, contain many individuals with slightly differing multi-locus SNP genotypes, inconsistent with the expectations for meiosis as the sole driver of genetic diversity ^19–21,23,30^.

The extreme plasticity of Phytophthora is well-known and although our data does not elucidate the processes driving DEA; knowing it exists is crucial. Phytophthora capsici produces millions of spores on a single vegetable fruit and the opportunities for adaptive and nearly instantaneous evolution are impressive. The challenges to implementing routine genetic analyses and constructing stable community resources are daunting. Although there are no other oomycetes with detailed linkage maps, it is important to recognize the importance of allele dosage when characterizing populations and conducting research. Although this work focuses on DEA as it manifests during zoosporogenesis, we’ve observed many instances where significant changes in pathogenesis and mating type occur spontaneously within individual cultures (without any obvious intervening zoospore formation) (Kurt Lamour, unpublished data). As a previous doctoral student of K. Lamour once lamented, “It is like trying to hold water in your hand and it just won’t sit still”.

## Online Methods

### Isolates, zoospore progeny, mating type and mefenoxam sensitivity

Field isolates were recovered using standard techniques where a small section of infected plant tissue is plated onto agar media amended with antibiotics and antifungal compounds as previously described ^7^. The recovery of single zoospore progeny is as previously described where agar plates are flooded, the plates briefly chilled and resulting swimming spores are induced to encyst and allowed to germinate on water agar prior to retrieval under a light microscope using a needle ^7^. Mating type and sensitivity to mefenoxam were assessed as previously described using known A1 and A2 mating types and percentage growth of an isolate on amended media compared to growth on control media, respectively ^24^. Release of swimming zoospores from the store-bought infected cucumber fruit (Figure 2) was induced by submerging the whole fruit in 500ml of sterile water and incubating for one hour at room temperature. The number of swimming spores was estimated by averaging 40 hemocytometer counts of 10ul aliquots of encysted zoospores.

### DNA, sequencing, analysis and data availability

Isolates were grown in nutrient broth, the resulting mycelium was freeze dried and powdered and genomic DNA extracted as described previously ^31^. The genomic DNA was sheared using a sonicator and PCR-free libraries constructed as previously described ^24^. The raw data was trimmed for quality, mapped to the *P. capsici* ordered (or unordered) genome, heterozygous allele frequencies assigned requiring >20X coverage and histograms constructed as previously described ^24^. Sequences for all isolates described in the paper are deposited in the NCBI under PRJNA386483.

---

Supplementary Figure showing ploidy by linkage group for 35 isolates of Phytophthora capsici or near relatives.

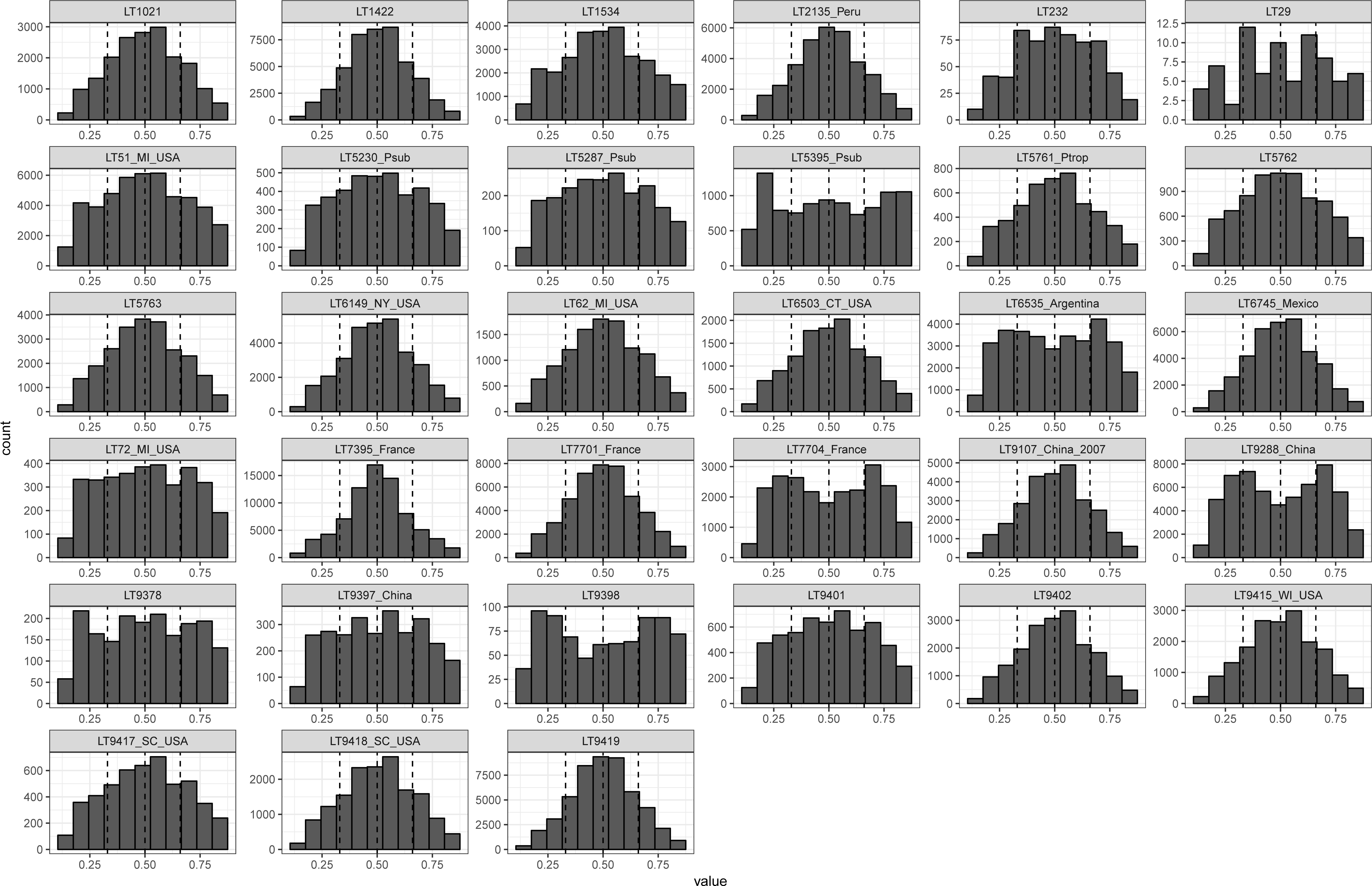

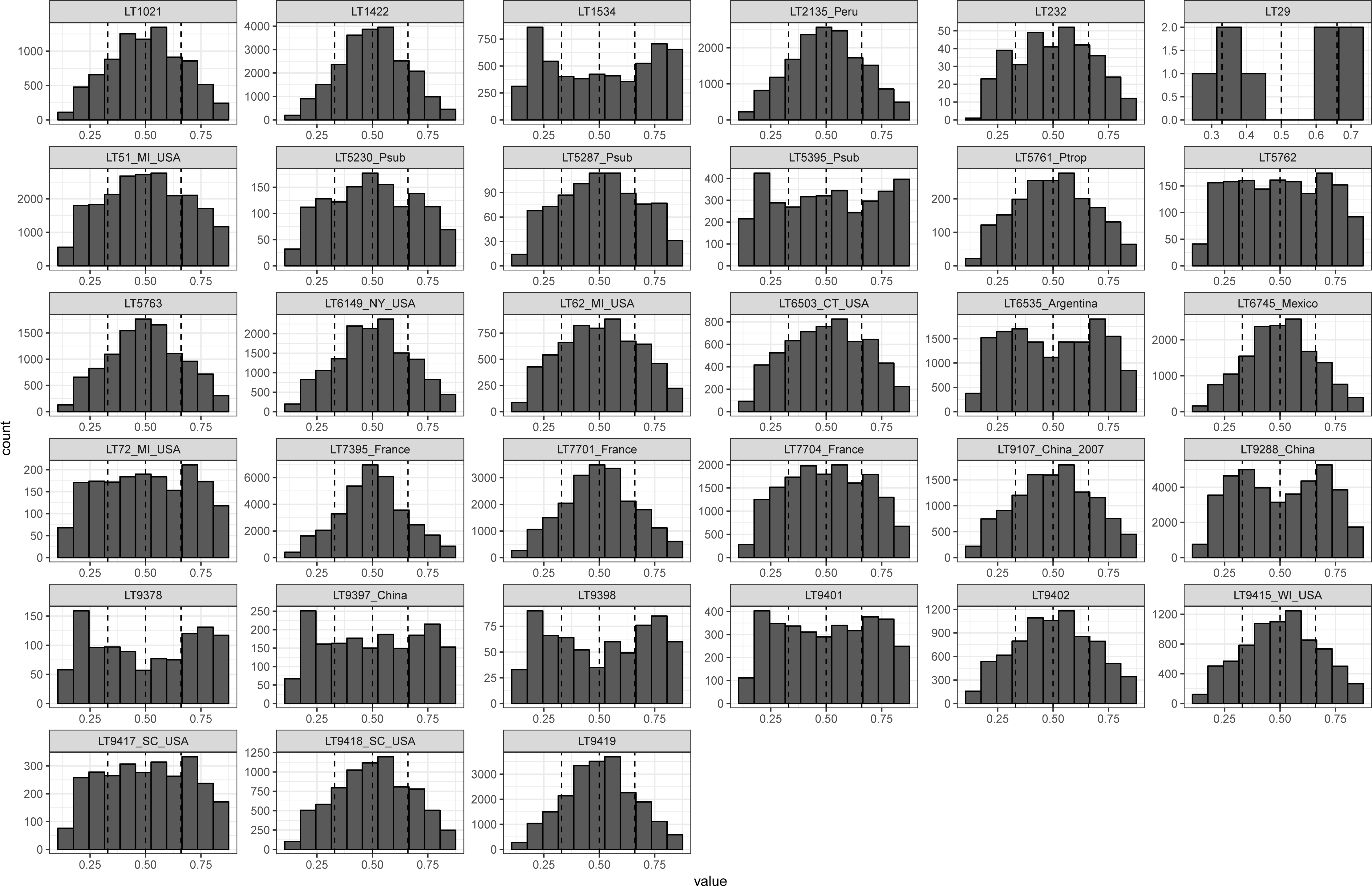

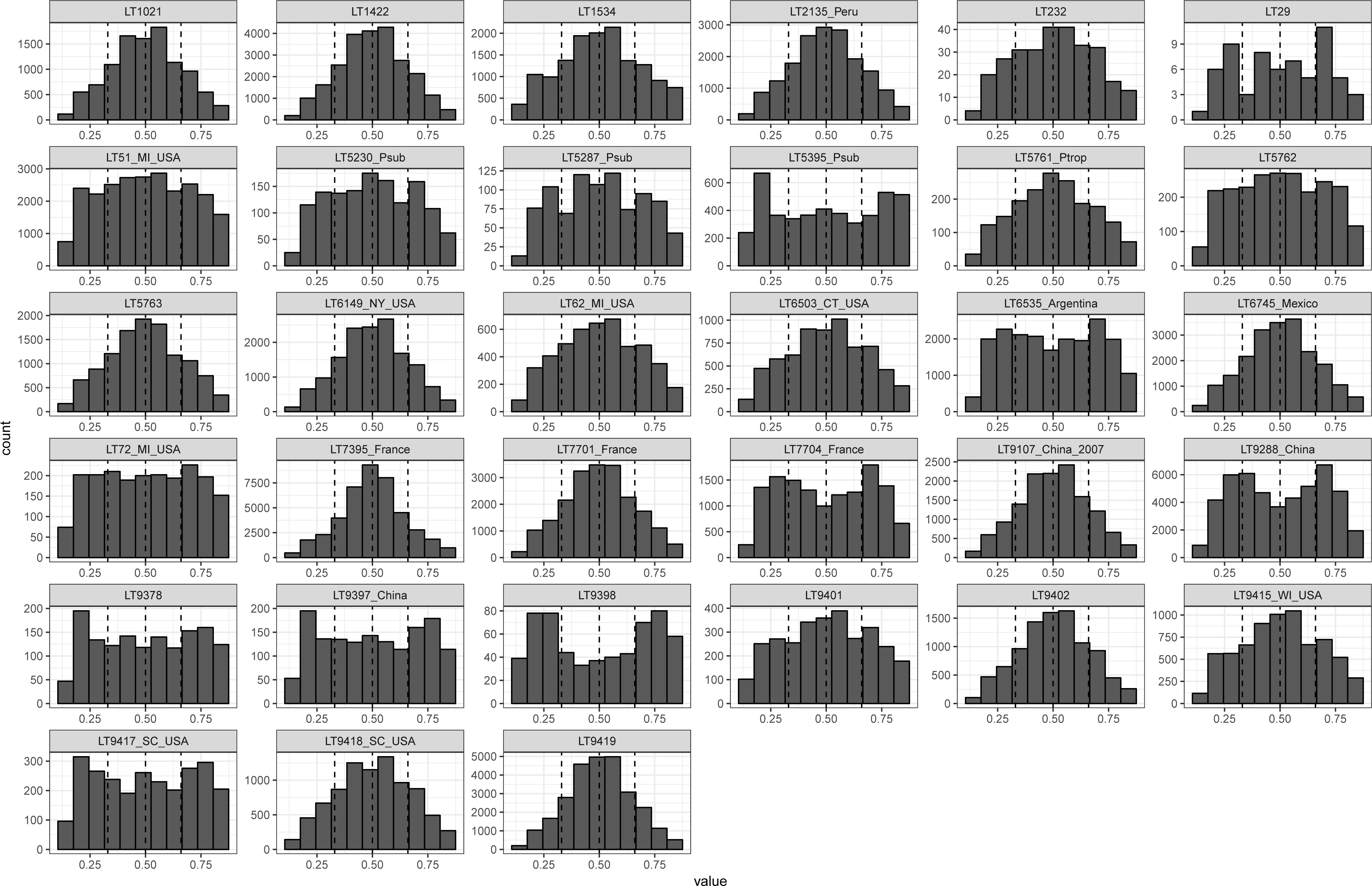

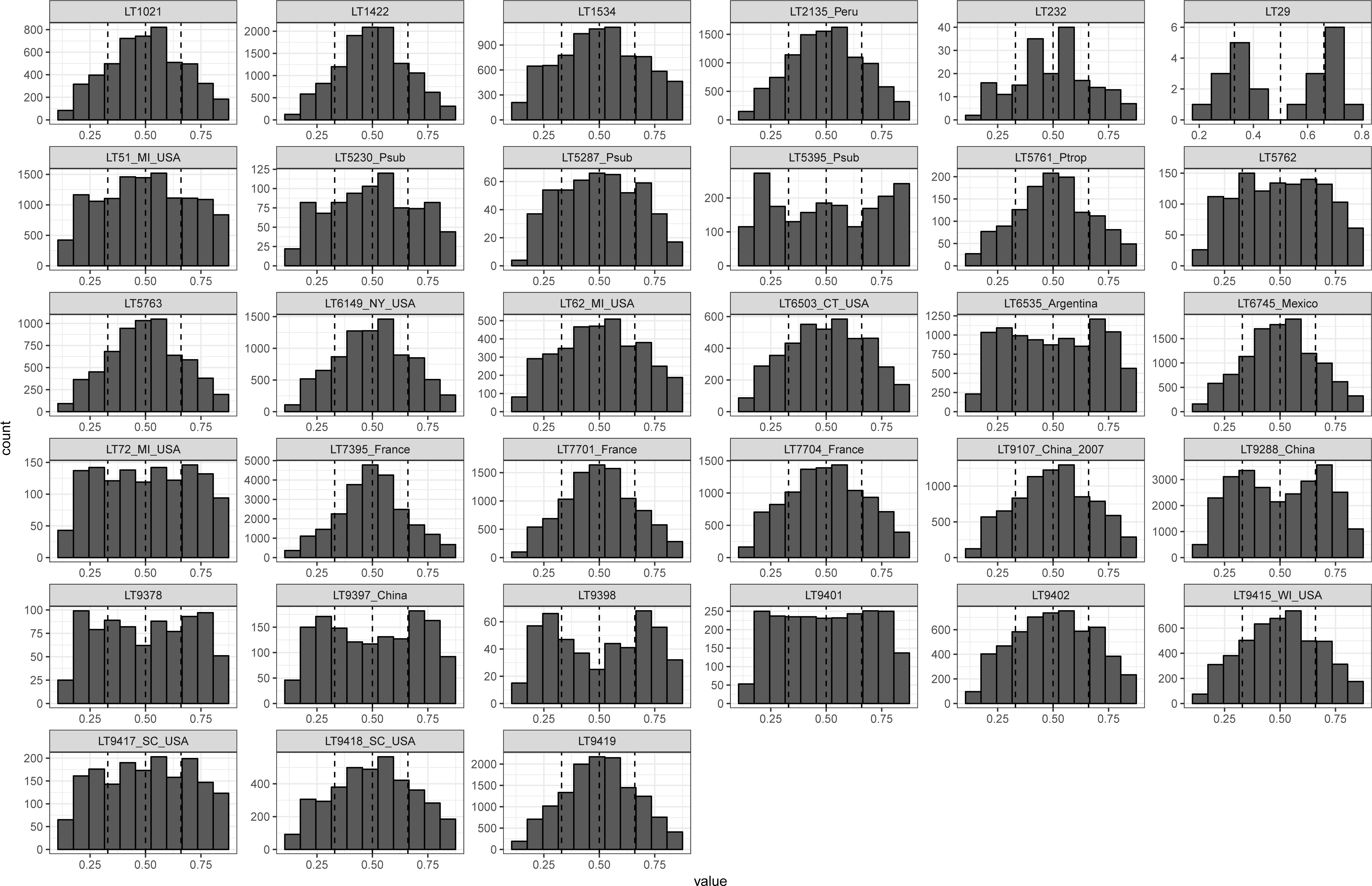

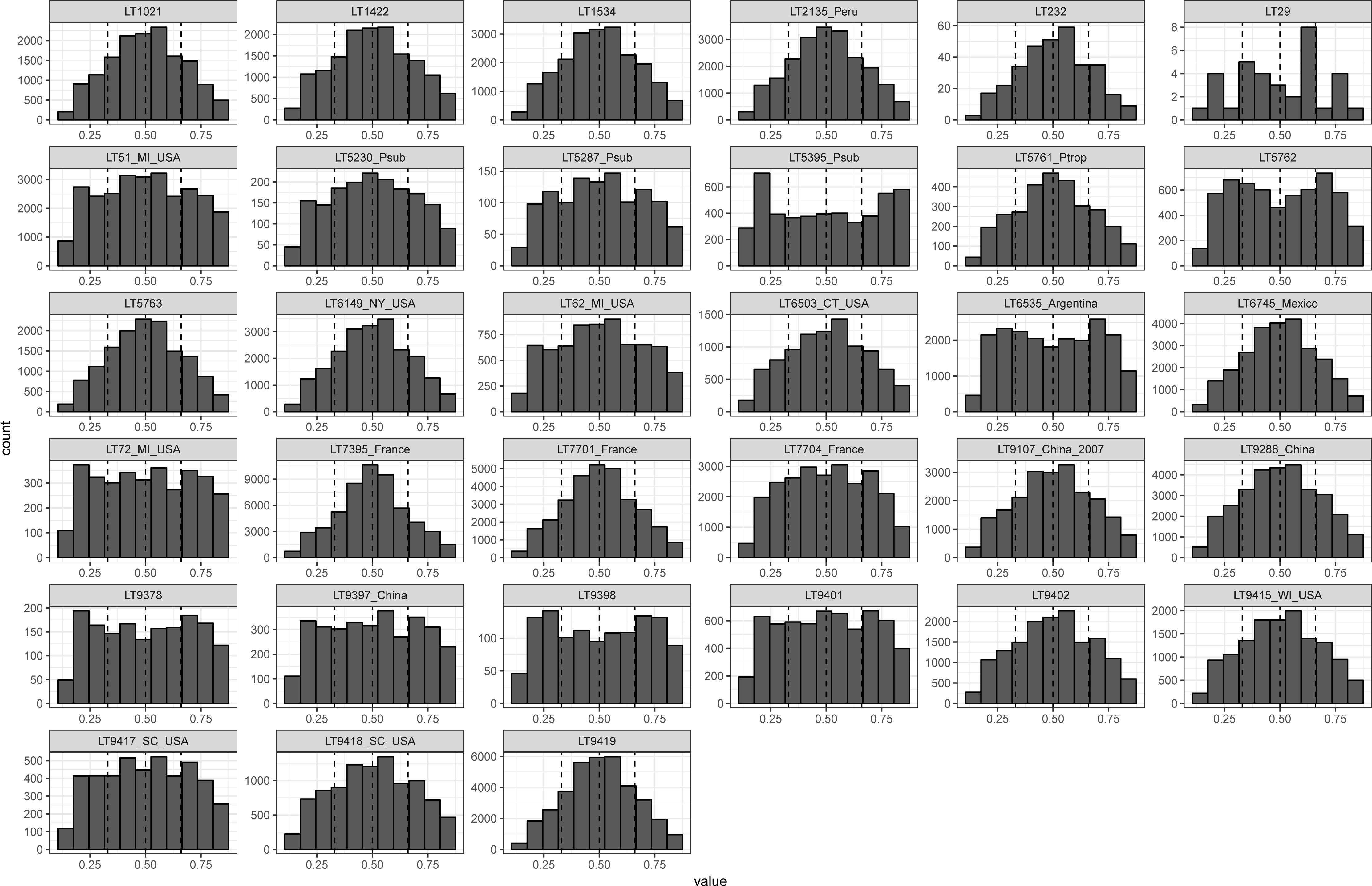

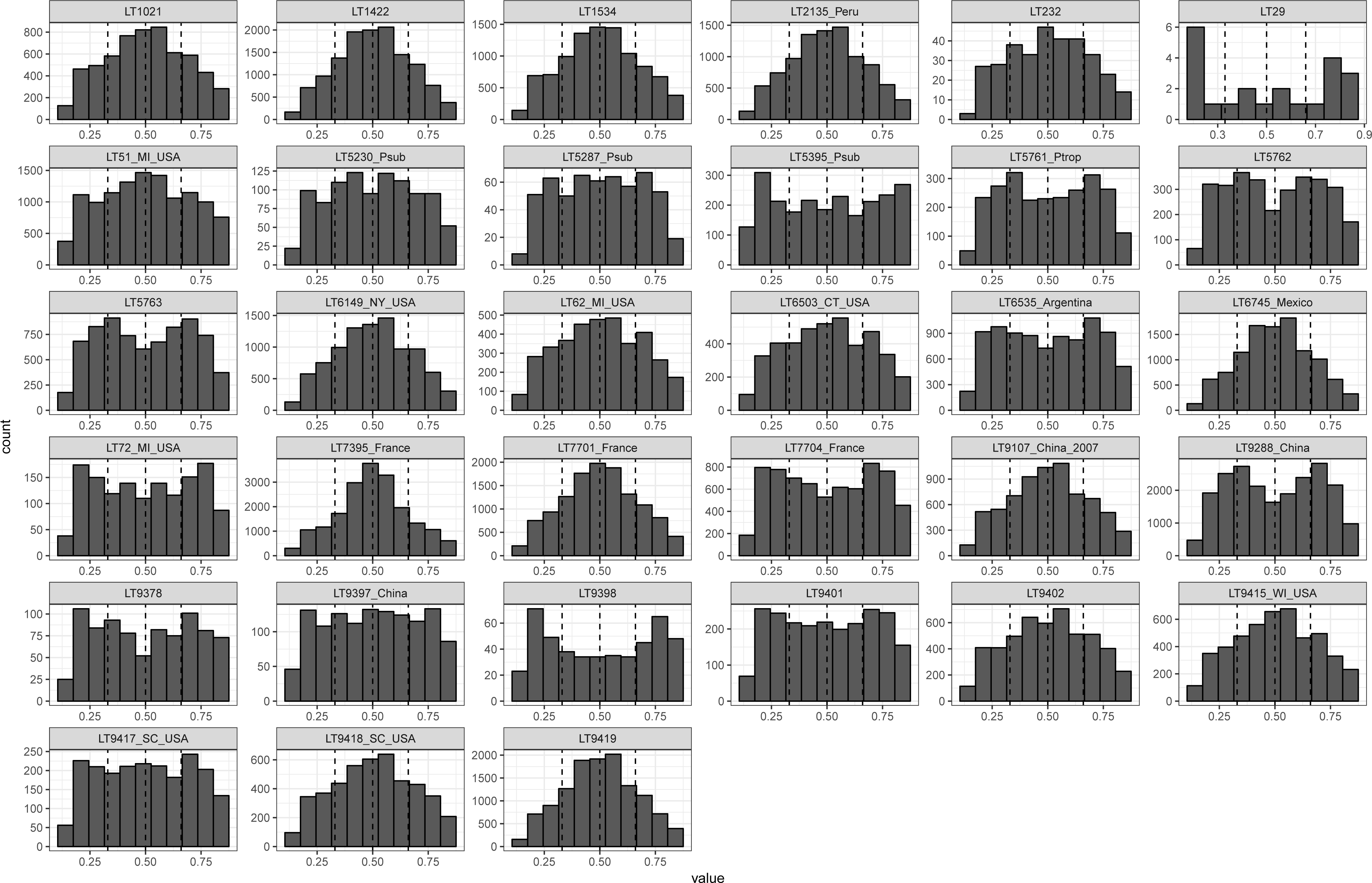

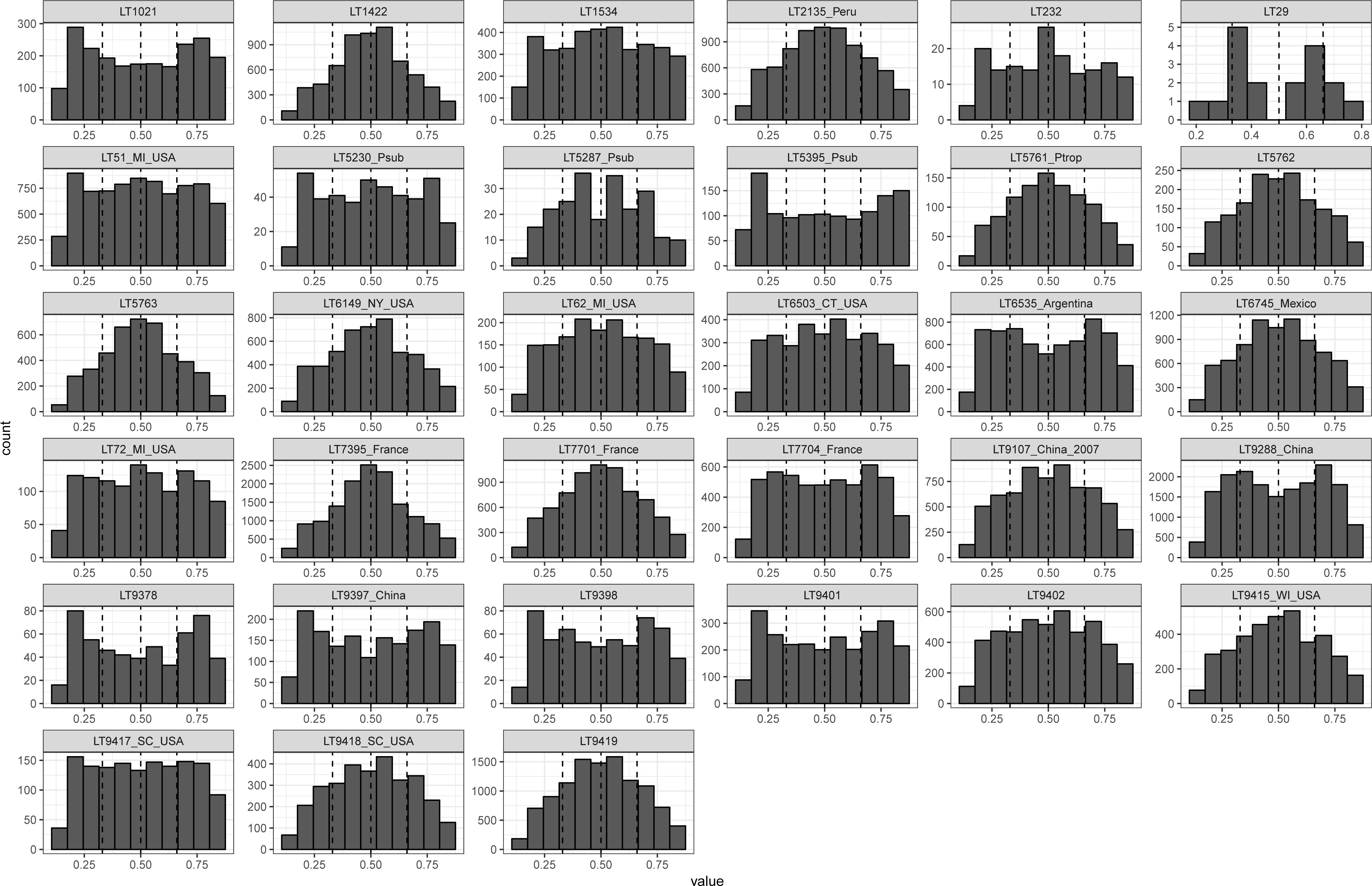

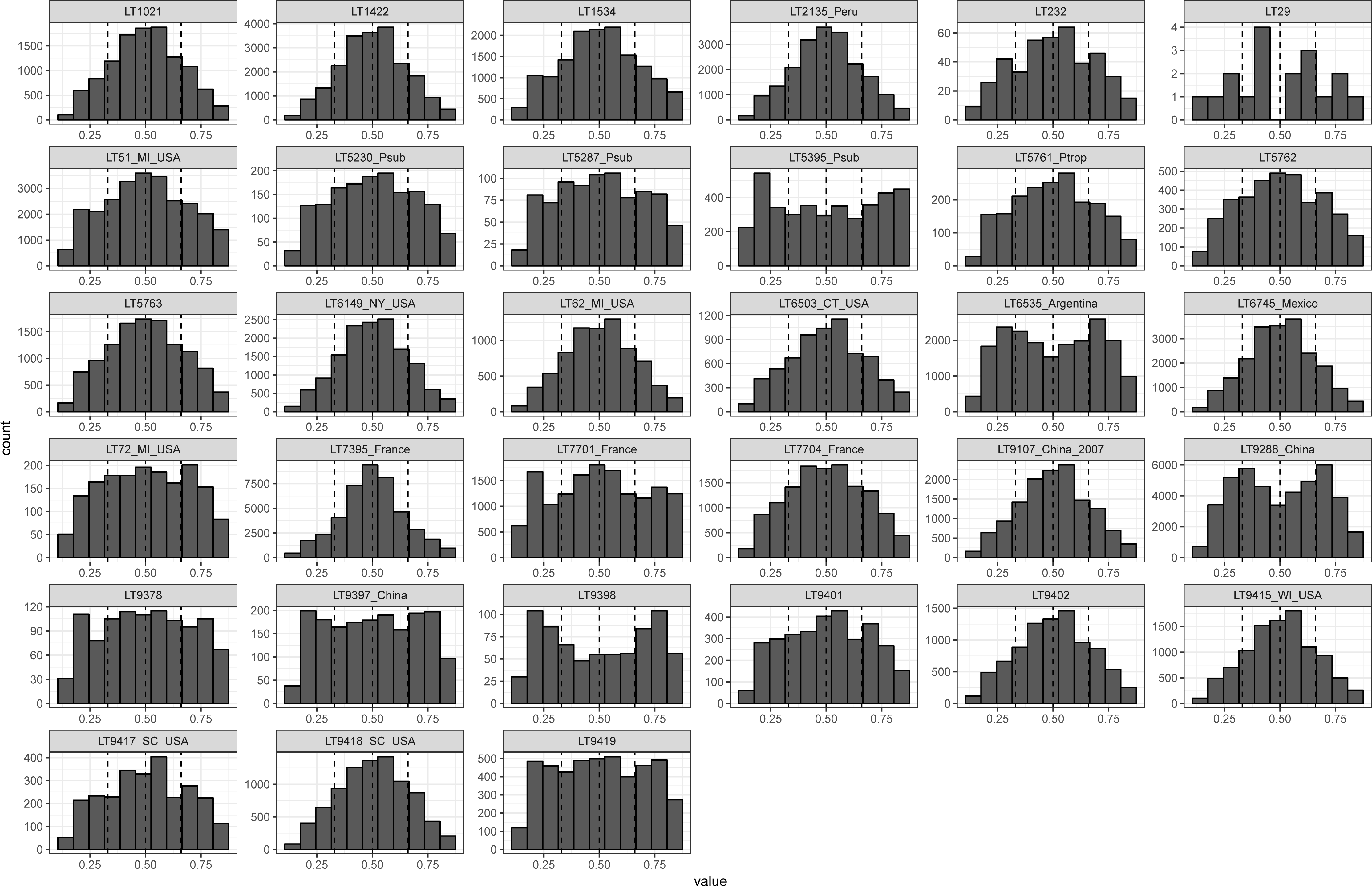

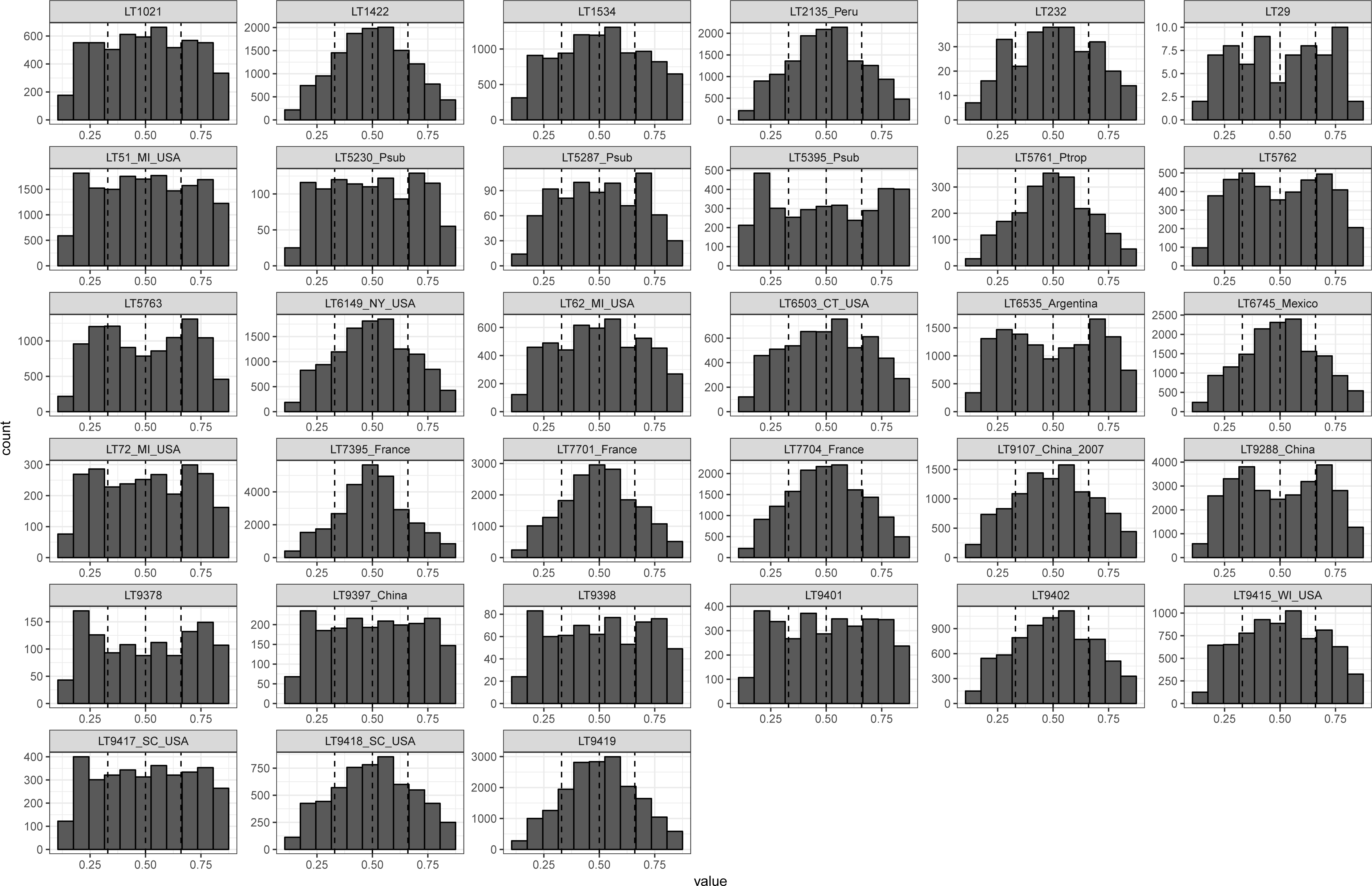

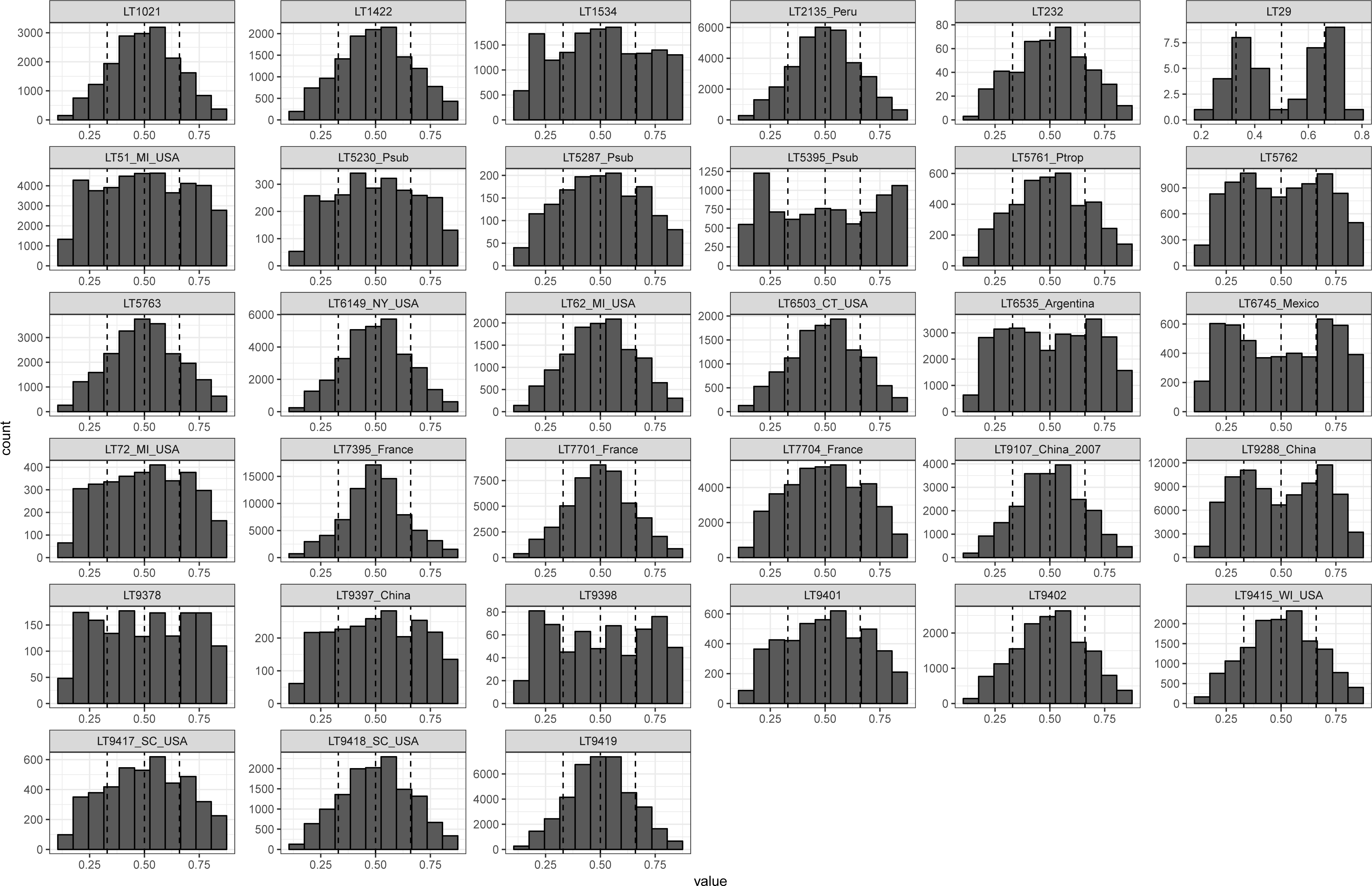

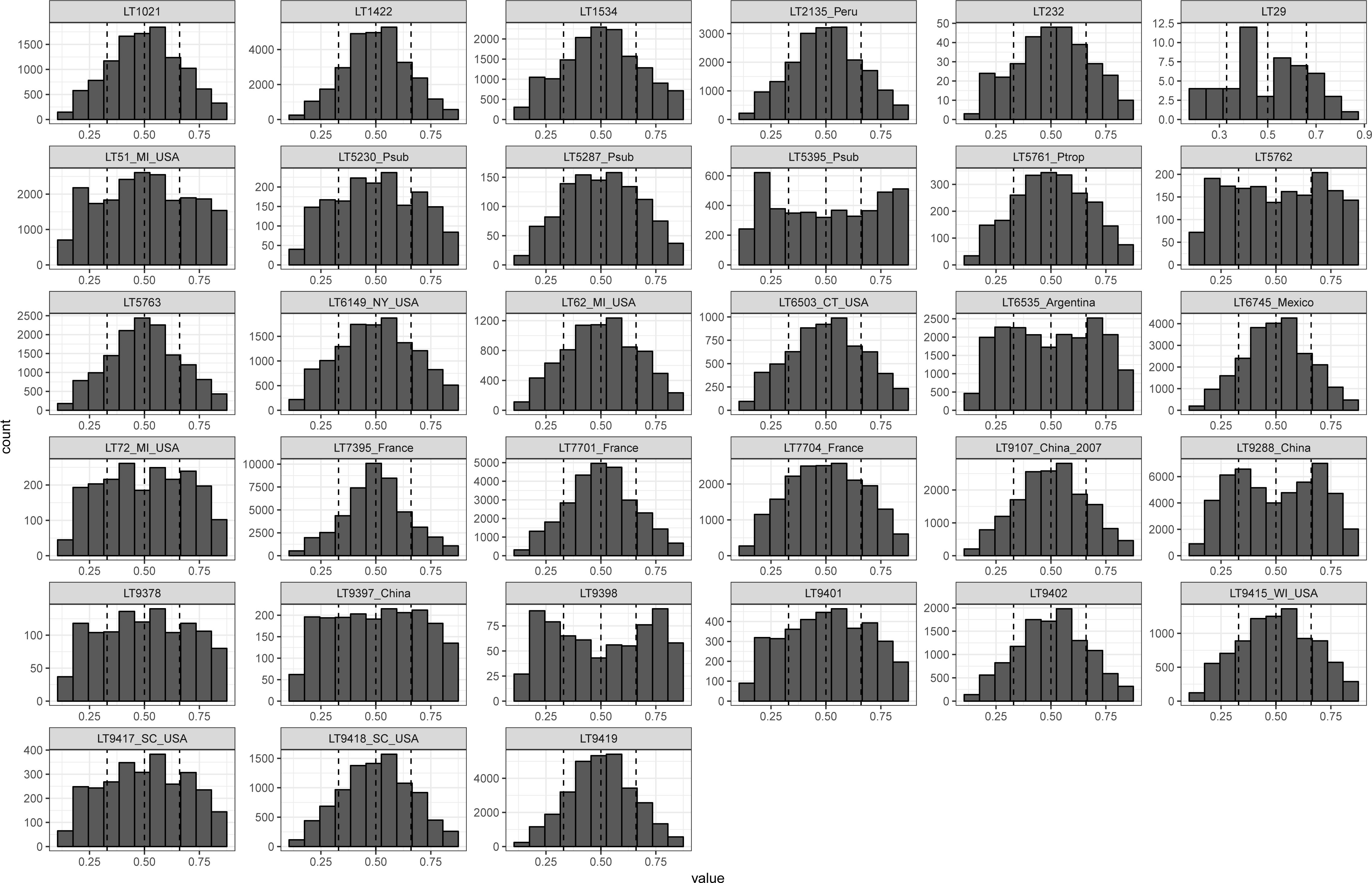

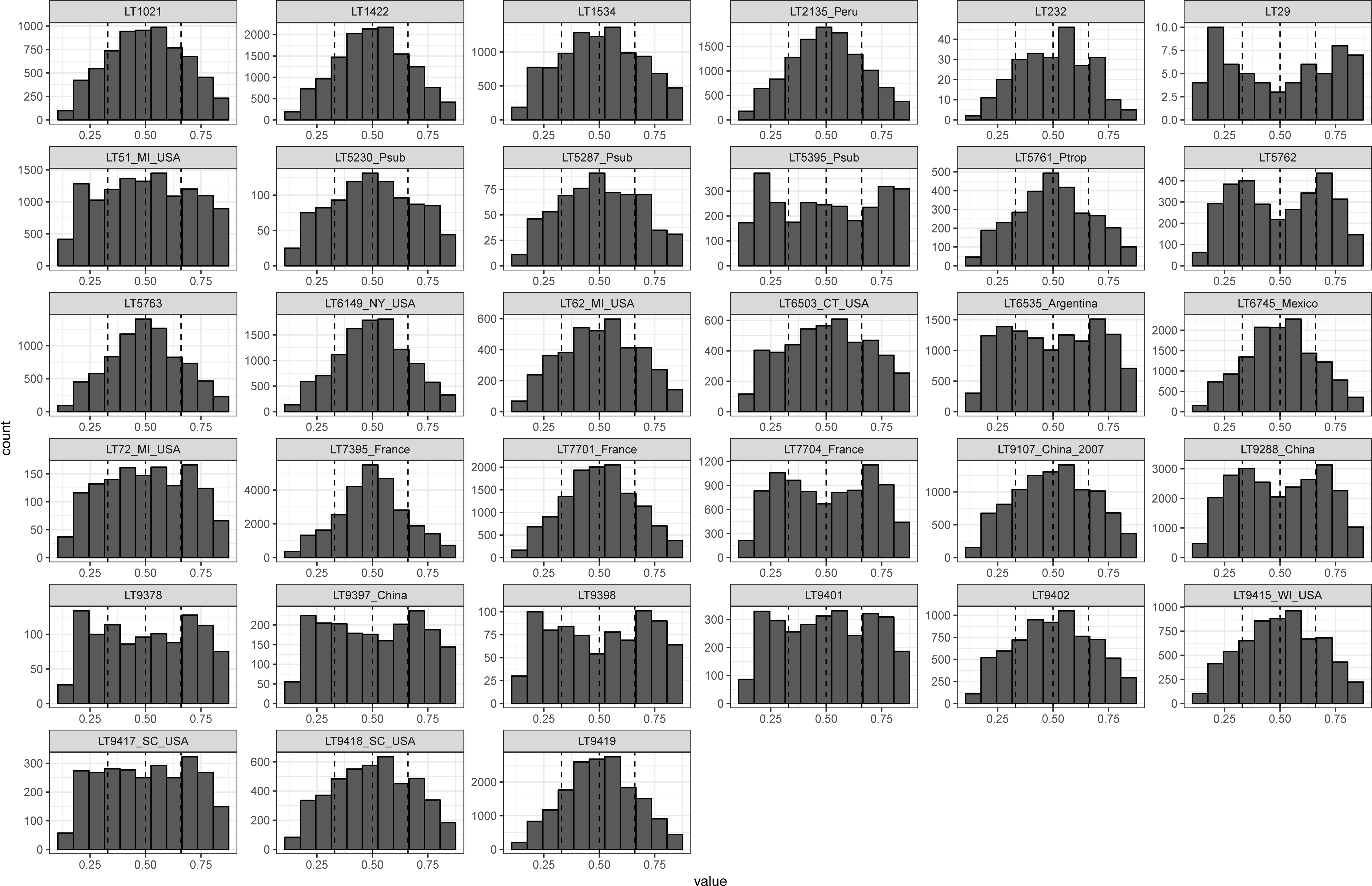

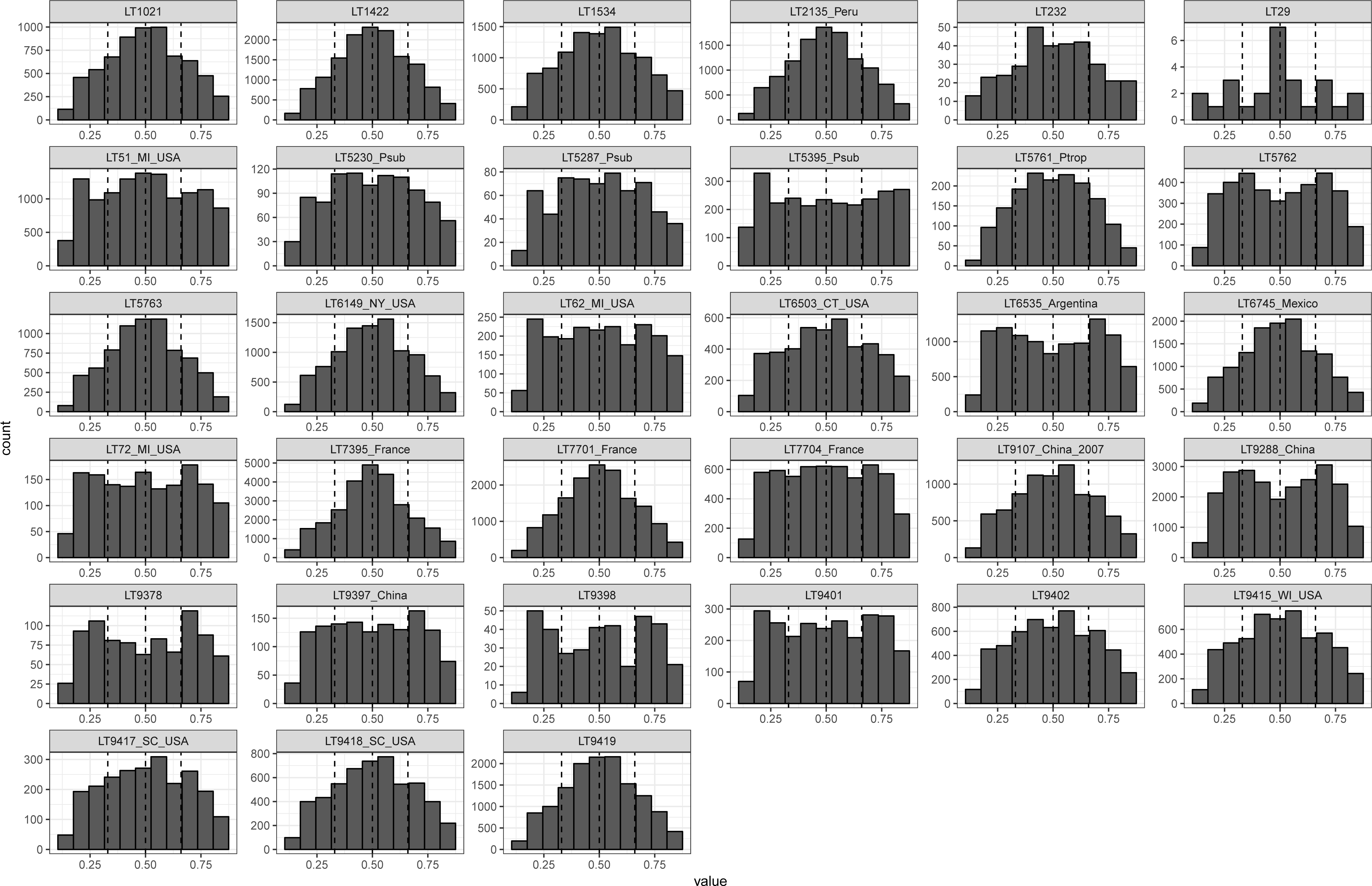

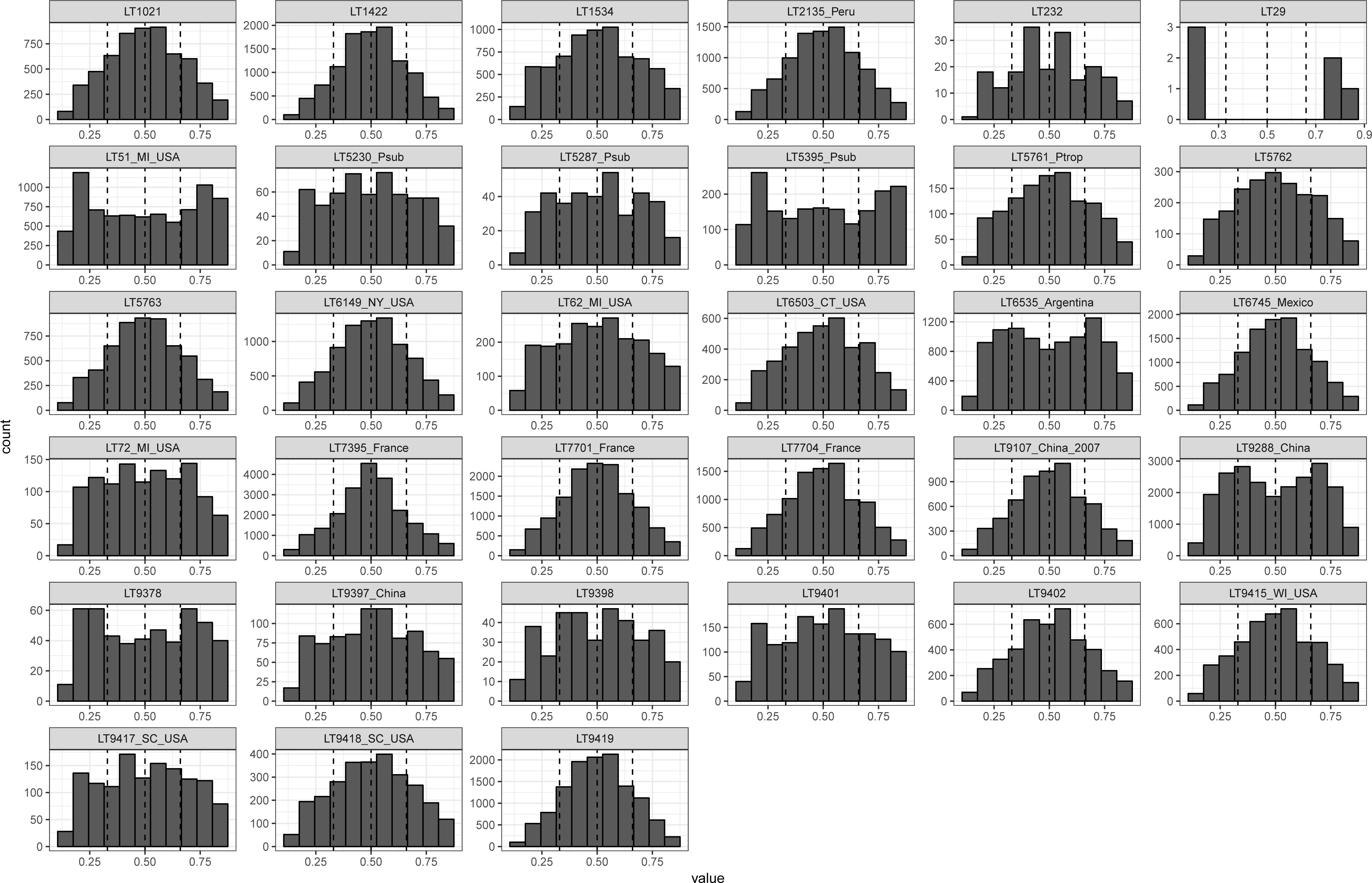

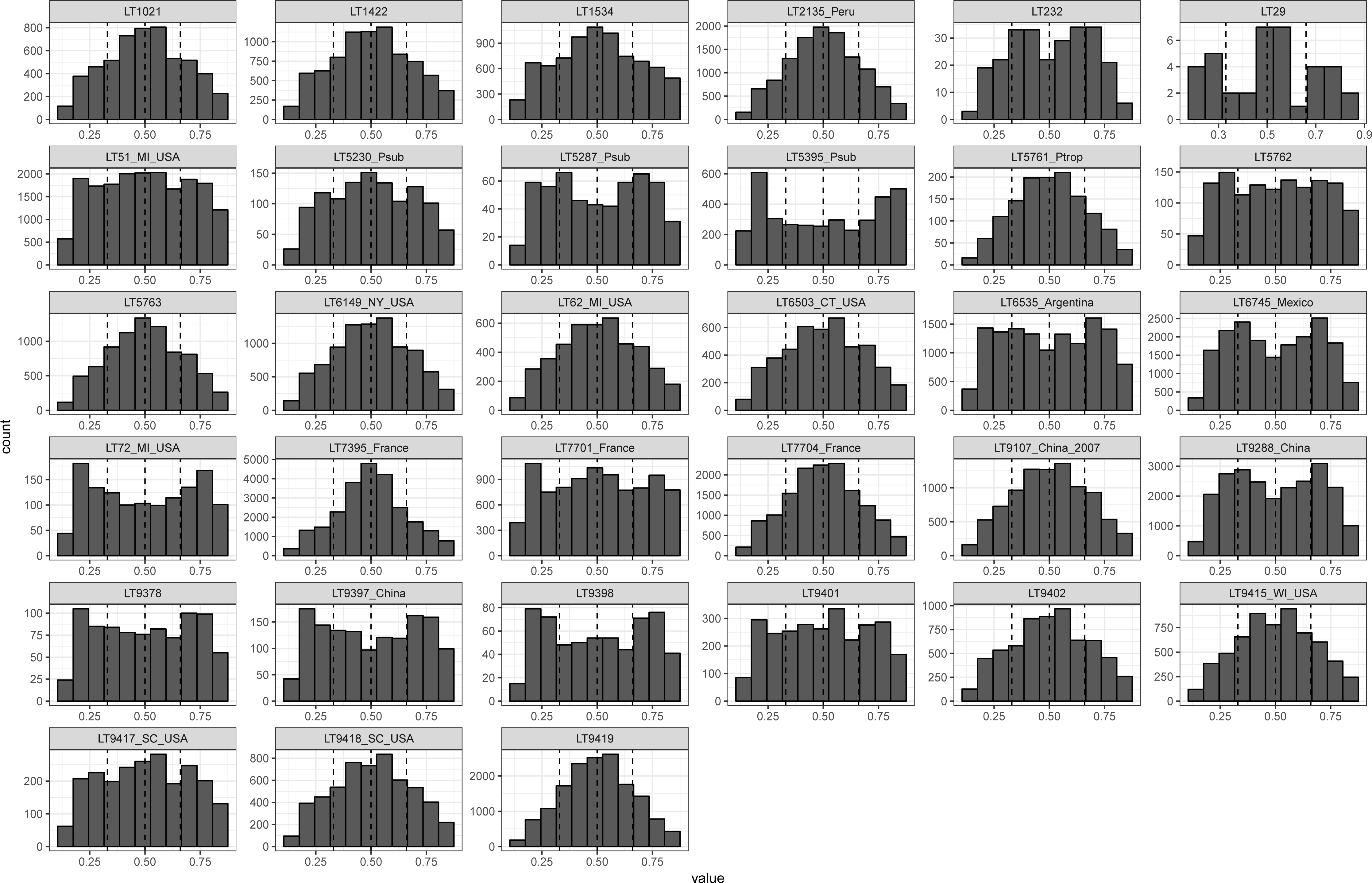

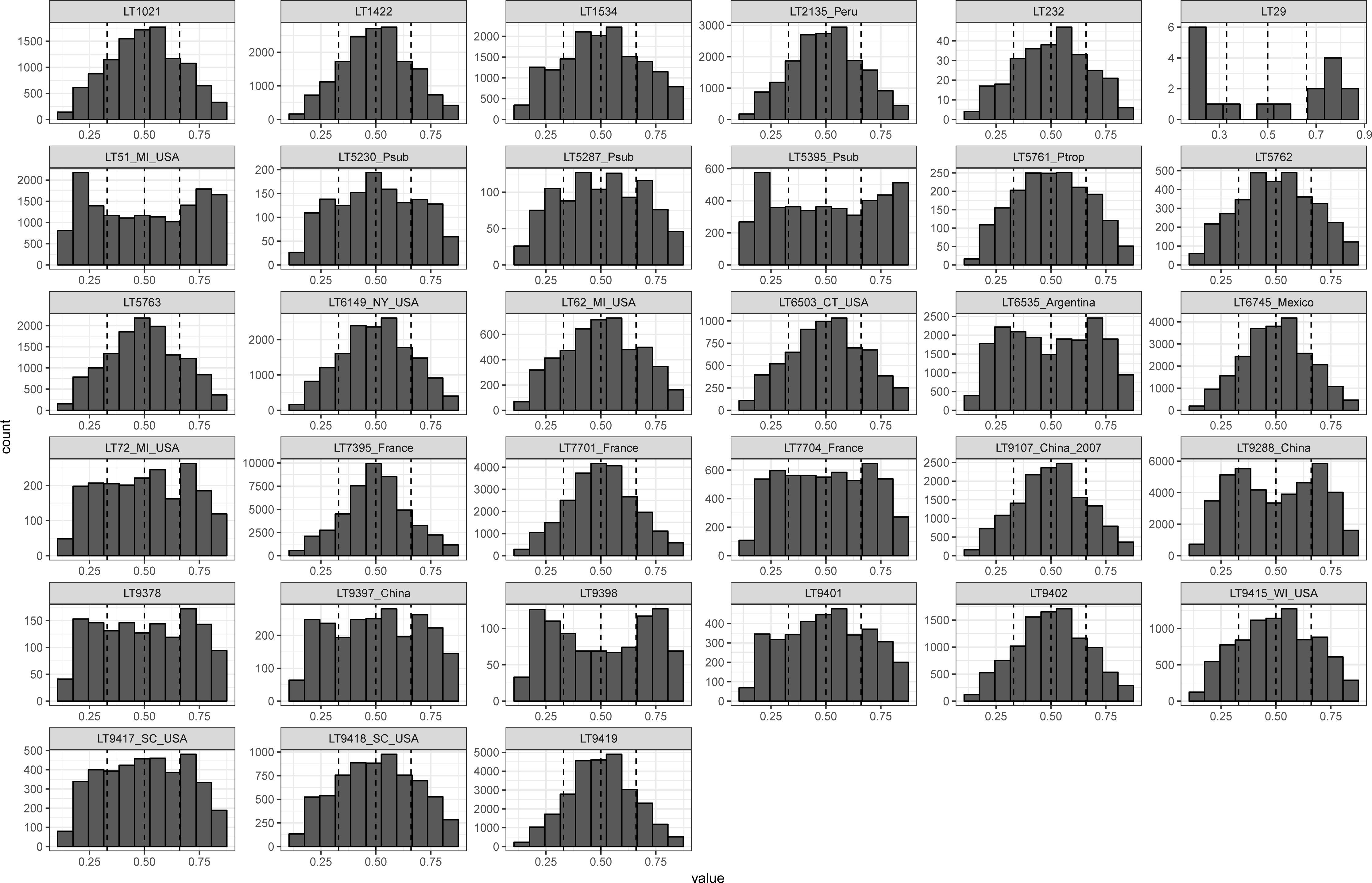

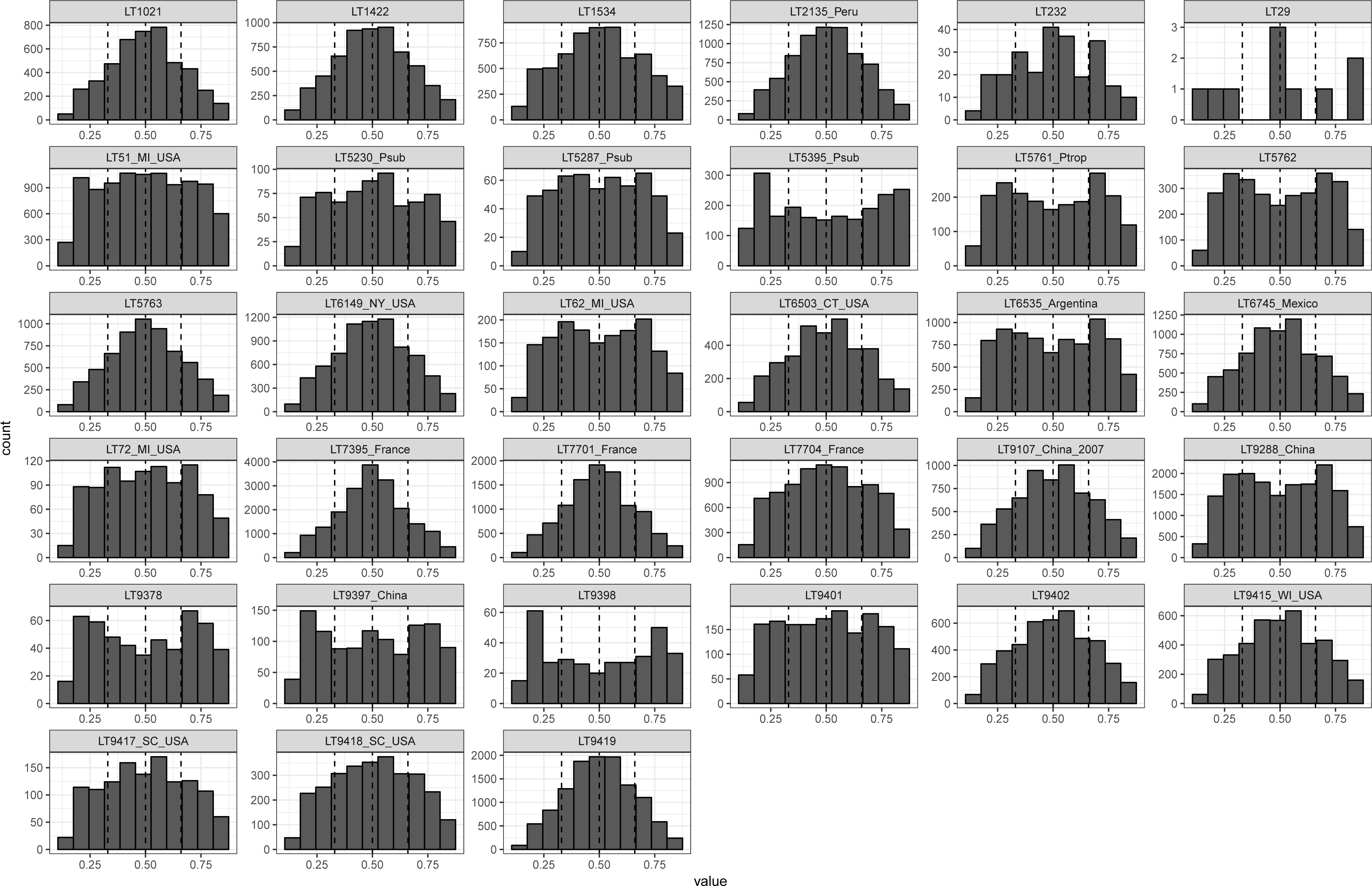

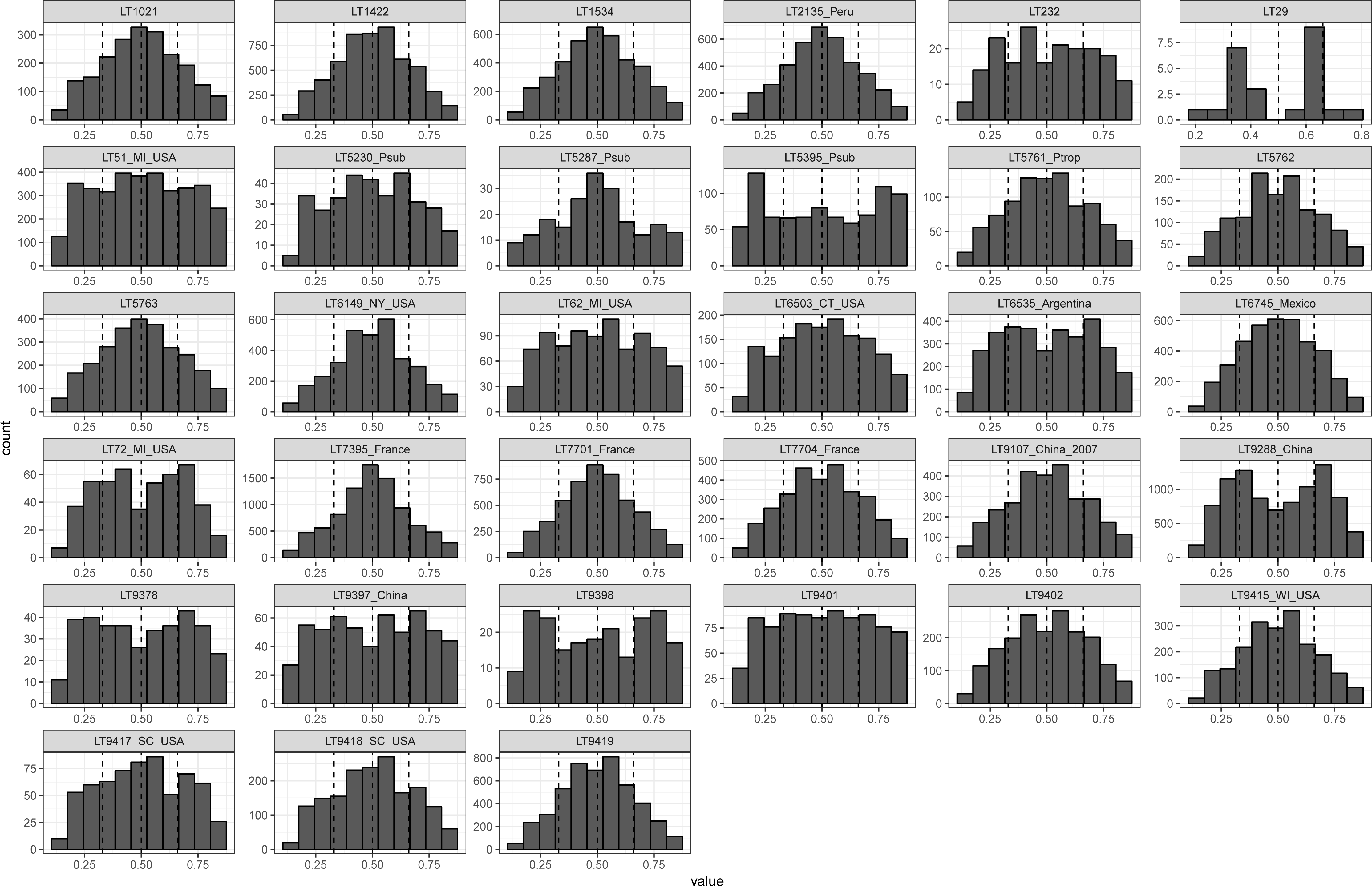

